# Insights into the aetiology of snoring from observational and genetic investigations in the UK Biobank (n=408,317)

**DOI:** 10.1101/808691

**Authors:** Adrián I. Campos, Luis M. García-Marín, Enda M. Byrne, Nicholas G. Martin, Gabriel Cuéllar-Partida, Miguel E. Rentería

**Affiliations:** Genetic Epidemiology Lab, QIMR Berghofer Medical Research Institute, Brisbane QLD Australia; Faculty of Medicine, The University of Queensland, Brisbane QLD Australia; Tecnológico de Monterrey, Escuela de Ingeniería y Ciencias, Zapopan, Jalisco, México; Institute for Molecular Bioscience, The University of Queensland, Brisbane, QLD, 4072, Australia; University of Queensland Diamantina Institute, Brisbane QLD Australia

## Abstract

We conducted the largest study of snoring using data from the UK Biobank (*n* ∼ 408,000; snorers ∼152,000). A genome-wide association study (GWAS) identified 42 genome-wide significant loci, with a SNP-based heritability estimate of ∼10% on the liability scale. Genetic correlations with body mass index, alcohol intake, smoking, schizophrenia, anorexia nervosa and neuroticism were observed. Gene-based associations identified 173 genes, including *DLEU7, MSRB3* and *POC5* highlighting genes expressed in brain, cerebellum, lungs, blood, and oesophagus tissues. We used polygenic scores (PGS) to predict recent snoring and probable obstructive sleep apnoea (OSA) in an independent Australian sample (n∼8,000). Mendelian randomisation analyses provided evidence that larger whole body fat mass causes snoring. Altogether, our results uncover new insights into the aetiology of snoring as a complex sleep-related trait and its role in health and disease beyond being a cardinal symptom of OSA.

## INTRODUCTION

Snoring is the vibration of upper airway structures that occurs when the muscles of the airway relax during sleep, thus creating noise as the air passes in and out while breathing. Habitual snoring is common in the population, its overall prevalence increases with age and is higher in males (35-45%) than females (15-28%).^1^ Importantly, snoring is a hallmark of OSA, a sleep-related breathing disorder characterised by repeated episodes of complete or partial obstructions of the upper airway during sleep, despite the effort to breathe.^1^ OSA is usually associated with a reduction in blood oxygen saturation, and is often accompanied by associated daytime symptoms, such as excessive daytime sleepiness, fatigue and decreased cognitive function. While the vast majority of patients with OSA exhibit snoring, a minority (20-25%) of patients with central sleep apnoea do not snore,^2^ and it is estimated that sleep apnoea may occur in as many as 20 to 40 percent of the adult population that are snorers, leaving the remaining 60-80% of snorers in the category of habitual non-apnoeic benign snorers. Snoring has previously been associated with body mass index (BMI)^3,4^ as well as with risk of cardiovascular disease, such as coronary heart disease and stroke among postmenopausal women.^5^ Twin and family studies have demonstrated the existence of a genetic predisposition to habitual snoring, with heritability estimates suggesting that 18-28% of variance can be accounted for by genetic factors^4,6^. A proportion of its heritability may be mediated through other heritable lifestyle factors such as smoking and alcohol consumption which can also contribute to snoring.^7–9^

Snoring is known to reduce sleep quality for both snorers and their sleeping partners,^10,11^ reducing energy/vitality and increasing daytime anxiety,^12^ risk of depression, stress, fatigue and sleepiness.^11^ Here, we leverage data from the UK Biobank and an Australian sample of adults, in an effort to characterise the molecular underpinnings of habitual snoring as a complex, polygenic trait, and investigate its relationship to known correlates such as BMI, smoking and alcohol consumption. We also employ statistical genetics methods such as LD score regression and Mendelian randomisation to identify genetically correlated traits with snoring and assess causality. Furthermore, we used our GWAS results to estimate individual PGS and predict snoring and probable OSA in an independent sample of Australian adults.

## RESULTS

### Snoring prevalence and risk factors

Our population-based discovery sample consisted of 408,317 individuals of white British ancestry from the UK Biobank. Participants in the sample were deemed as snoring ‘cases’ (37%) based on their report that a partner or housemate had complained to the participant about their snoring (see <u>Methods</u> and Table 1). Snoring was significantly associated with age (OR=1.011 [per year, 95% CI 1.009-1.012]) and, to a greater extent, with sex (OR_males_=2.264 [2.212-2.316]). The prevalence of sleep apnoea was higher within the snorer group (Table 1). Furthermore, BMI, smoking frequency and alcohol consumption frequency were also associated with snoring (Figure 1a). While snoring prevalence was higher in males, BMI was positively correlated with snoring prevalence in both males and females (Figure 1b). Smoking frequency was positively correlated with snoring prevalence in females, and to a lesser extent in males (Figures 1a and 1c). In contrast, alcohol consumption frequency was correlated with snoring in males, and to a lesser extent in females (Figures 1a and 1d). We further identified other factors such as whole body fat mass and sleep duration that are correlated with snoring (Supplementary Table 1).

**Table 1.**
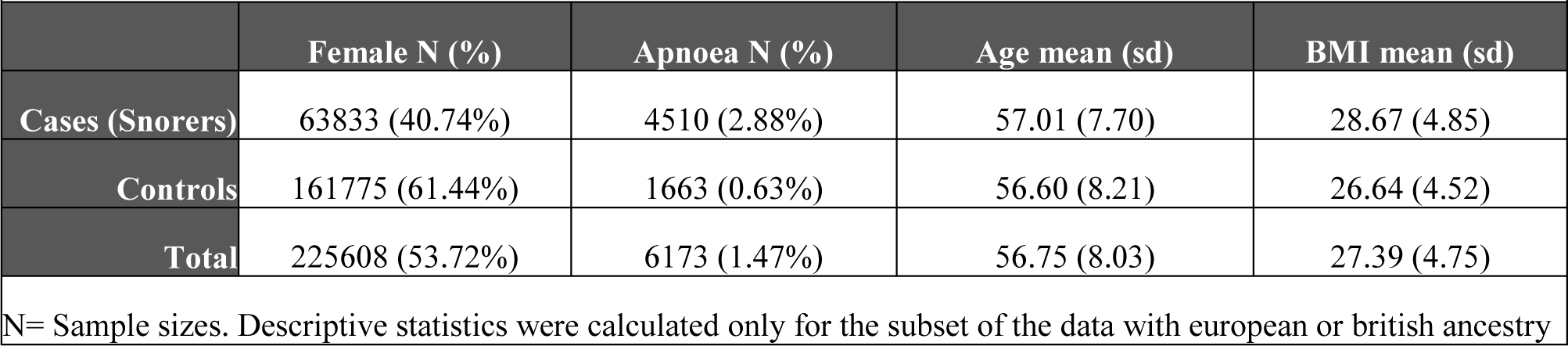
Sample composition and descriptive statistics of UK Biobank discovery sample

|  |  |  |  |  |
| --- | --- | --- | --- | --- |
|  | **Female N (%)** | **Apnoea N (%)** | **Age mean (sd)** | **BMI mean (sd)** |
| **Cases (Snorers)** | 63833 (40.74%) | 4510 (2.88%) | 57.01 (7.70) | 28.67 (4.85) |
| **Controls** | 161775 (61.44%) | 1663 (0.63%) | 56.60 (8.21) | 26.64 (4.52) |
| **Total** | 225608 (53.72%) | 6173 (1.47%) | 56.75 (8.03) | 27.39 (4.75) |
| N= Sample sizes. Descriptive statistics were calculated only for the subset of the data with european or british ancestry |  |  |  |  |

**Figure 1.**
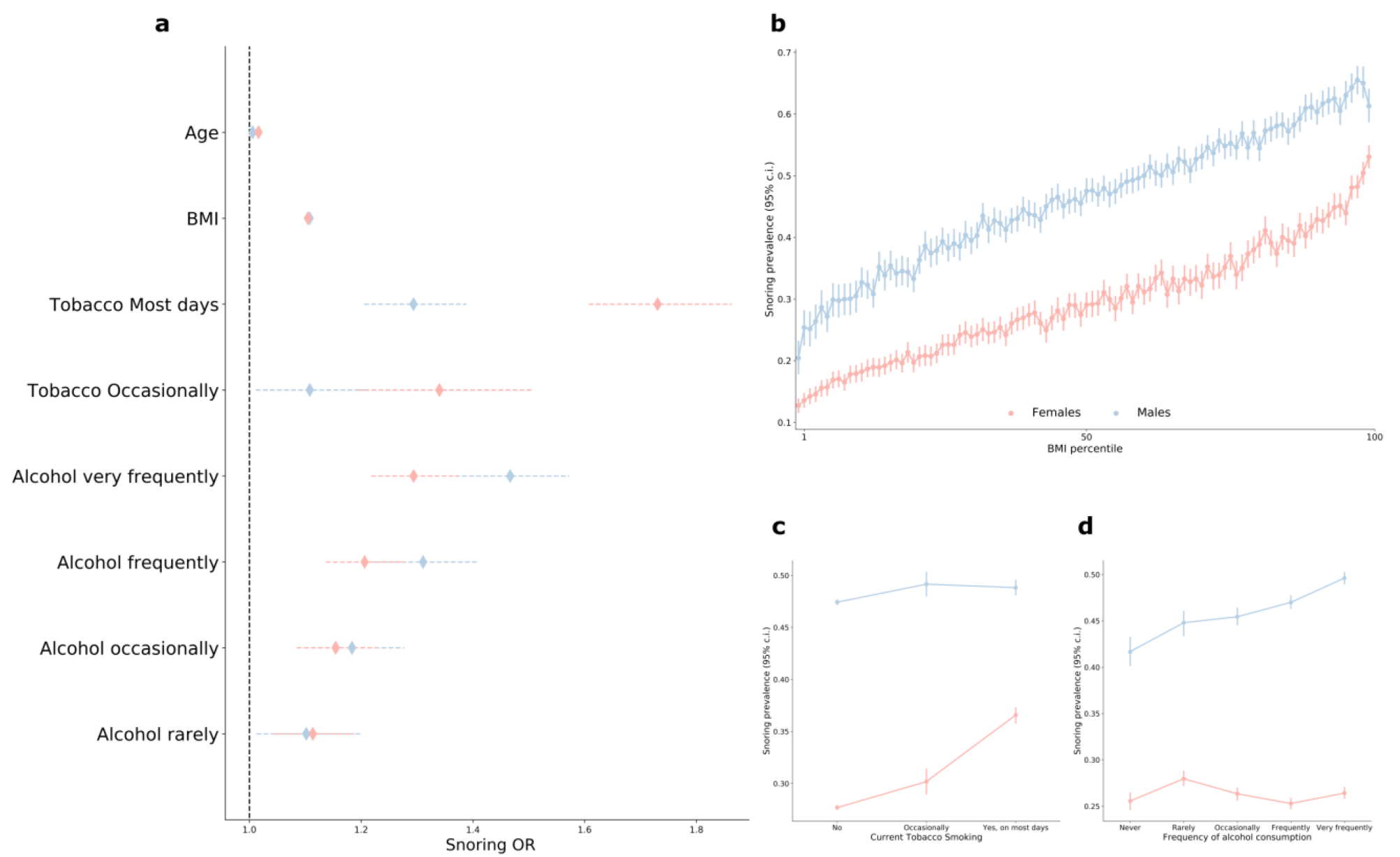
Sex, BMI, frequency of alcohol consumption and smoking are associated with increased snoring. a) Forest plot depicting the odds ratios of studied variables on snoring for males (blue) and females (red). b) Sex and BMI stratified prevalence of snoring. c) Sex and frequency of tobacco smoking stratified snoring prevalence. d) Sex and frequency of alcohol consumption-stratified snoring prevalence. Error bars denote the 95% confidence intervals estimated using the odds ratio and standard error (a) or 1000 pseudo replicates from a bootstrap resampling procedure (b, c and d)

### Discovery GWAS and SNP heritability

We performed a GWAS of snoring, taken as a dichotomous variable (*n* = 408,317; cases∼152,000; controls∼256,000). After quality control (see <u>Methods</u>), 11,010,159 genetic variants remained in the analysis. This uncovered 127 independent genome-wide significant associations across 41 genomic risk loci (Figure 2 and Supplementary Figure 1).^13^ Annotation for the top 15 risk loci is shown in Table 2, and a list of all genomic risk loci is given in Supplementary File 1. The overall SNP heritability on the liability scale (*h*^*2*^_*SNP*_) was 9.9% (S.E.=0.39%).

**Table 2.**
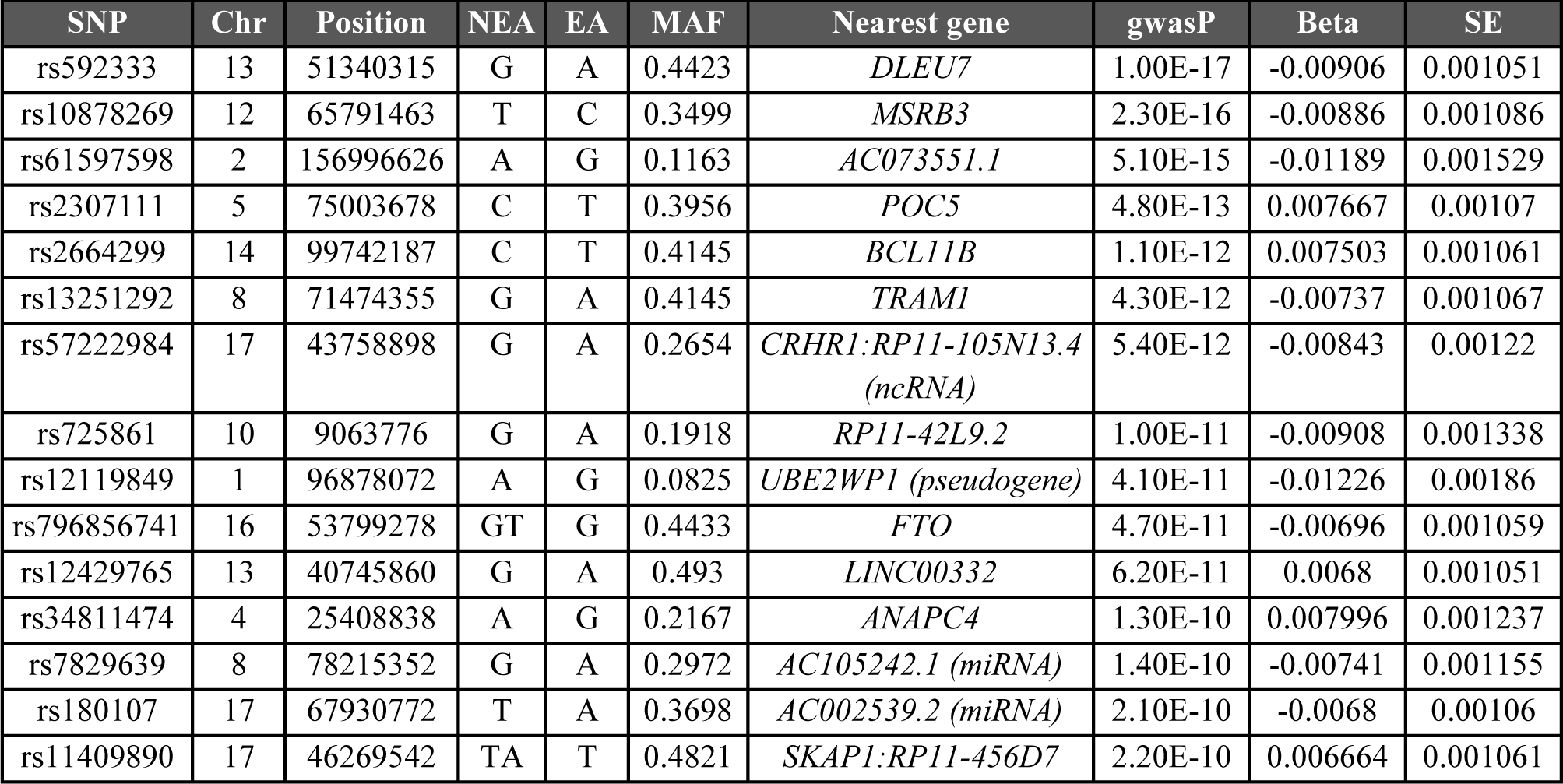
Top 15 genomic risk loci for snoring showing the top SNP for each locus.

**Figure 2.**
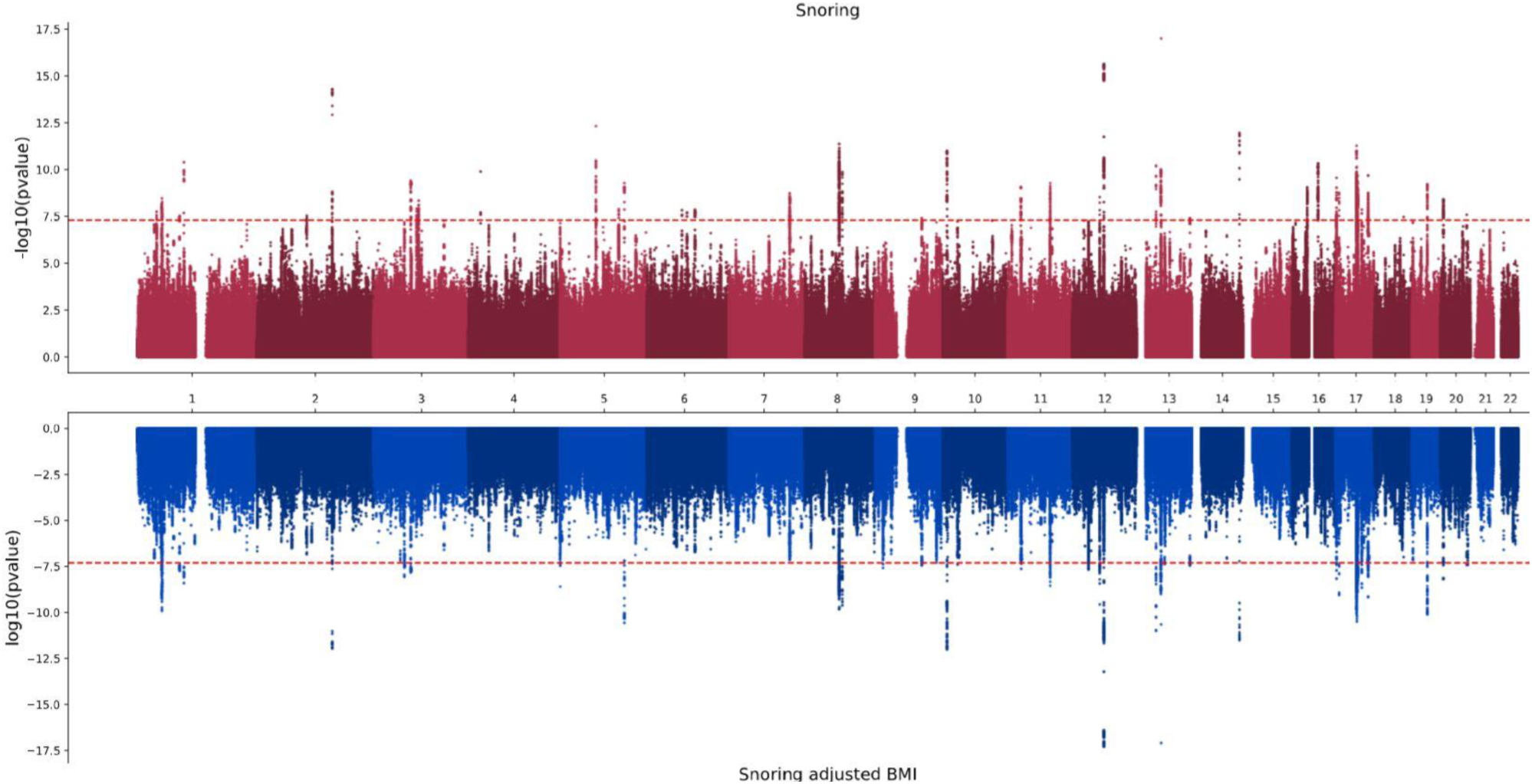
Genetic variants associated with snoring with (bottom) and without (top) adjustment for BMI. Results from the genome-wide association studies are presented as mirrored Manhattan plots. The x axis contains a point for each genetic variant passing QC and is ordered by chromosome and base position. The distance between each variant and the x-axis is a function of the significance (p-value) of the association. For the top panel the -log10(p-value) is graphed on the y_axis whereas the log10(p-value) is shown for the bottom panel. The red dotted line denotes the genome-wide significance threshold (p<5e-8).

### Genetic correlations

The trait that showed the highest genetic correlation with habitual snoring was self reported sleep apnoea (r_G_=0.78 S.E.=0.17, p-value=3×10^−05^; Supplementary File 2). Other genetically correlated traits with snoring included BMI, whole body fat mass, sodium in urine, mood swings, coronary artery disease, alcohol intake frequency, pulse rate, current tobacco smoking, heart disease, lung cancer, the ratio between forced expiratory volume in 1 second (FEV1) and forced vital capacity (FVC), neuroticism, subjective wellbeing and heart rate, among others. Traits showing a negative genetic correlation with snoring included schizophrenia, FVC, FEV1, fluid intelligence score, educational attainment, age at menarche, mean accumbens volume and anorexia nervosa. Overall, traits related to body-mass index, risk for psychiatric disease, lung function and heart disease were among those with the strongest evidence of association (Figure 3 and Supplementary File 2). Notably, pulse rate, whole body fat mass and BMI were also phenotypically associated with snoring in this sample (see above and Supplementary Table 1).

**Figure 3.**
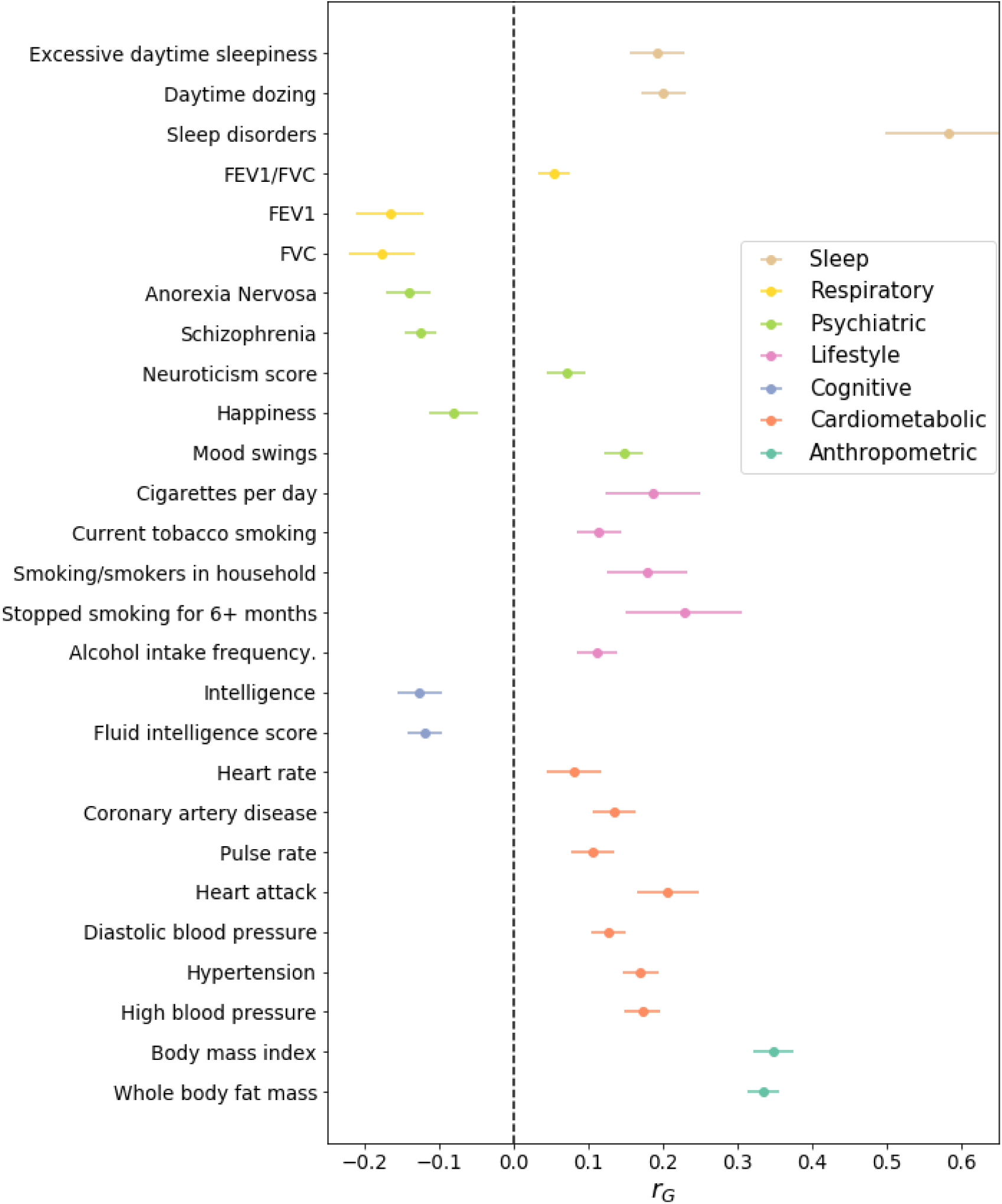
Snoring is genetically correlated with lifestyle, psychiatric, respiratory and other complex traits. Circles represent LD score based estimates of the genetic correlation between snoring and other complex traits (y-axis). All of the depicted traits had a statistically significant genetic correlation with snoring after multiple testing corrections (benjamini-hochberg; fdr <0.05). Error bars show the s.e. Not all significant associations are shown due to lack of space, complete results are available in Supplementary File 2.

### Sensitivity Analysis

We performed a follow-up sensitivity GWAS for snoring including BMI as a covariate to explore the effects of BMI on associated variants. The results revealed 97 genome-wide significant SNPs across 34 genomic risk loci (Figure 2) with overall SNP heritability on the liability scale (*h*^*2*^_*SNP*_) of 8.67% (S.E.=0.39) (Table 3). Traits that remained genetically correlated with snoring after adjusting for BMI included schizophrenia, educational attainment, sodium in urine and sleep related traits such as daytime dozing, sleep apnoea and excessive daytime sleepiness (Supplementary File 3). The genetic correlation between both the adjusted and unadjusted GWAS was high (*r*_G_=0.923, S.E.=0.003, p-value=1×10^−300^) suggesting that a considerable amount of snoring predisposition is not fully explained by BMI.

**Table 3.** SNP-based heritability of snoring on the liability scale.

| Trait | $h^2_{SNP}$ | SE | $\lambda_{GC}$ | intercept |
| --- | --- | --- | --- | --- |
| Snoring | 9.9% | 0.39% | 1.4281 | 1.0427 (0.011) |
| Snoring adj. for BMI | 8.67% | 0.39% | 1.3685 | 1.0314 (0.0094) |
| Snoring males | 8.77% | 0.54% | 1.2005 | 1.0164 (0.0078) |
| Snoring females | 12.42% | 0.57% | 1.254 | 1.021 (0.0074) |
| Snoring adj. for BMI males | 7.72% | 0.56% | 1.2005 | 1.017 (0.008) |
| Snoring adj. for BMI females | 10.85% | 0.54% | 1.2531 | 1.0237 (0.007) |

LD score regression derived SNP based heritability results. Estimates were transformed to the liability scale assuming equal population and sample prevalences. $\lambda_{GC}$ is the genomic inflation factor and intercept is the LD score regression intercept

### Positional, eQTL and gene-based test prioritisation

To gain insights into the functional consequences of individual genome-wide significant variants, we used positional and eQTL mapping as well as genome-wide gene-based association analyses. From positional and eQTL mapping, we identified 149 protein coding genes mapping to a genome-wide significant SNP. The nearest genes to the top signals included *DLEU7* on chromosome 13, and *MSRB3* on chromosome 12. In addition to *DLEU7* and *MSRB3*, other compelling genes (prioritised by positional or eQTL mapping) for snoring included *BCL11B, FTO, SMG6, ROBO2, NSUN3, SNAP91* and *BCL2*, which have previously been associated with smoking;^14,15^ *BLC11B, FTO*^*16*^, *RNA5SP471*^*16,17*^ and *SND1* and *NSUN3*, previously associated with alcohol consumption;^14,16–18^ *FTO* and *SND1*, associated with coffee consumption;^19^ *LMO4* associated with insomnia;^20^ and *RNA5SP471* with narcolepsy.^20,21^ Also, *ROBO2*, previously associated with chronotype;^22,23^ and multiple genes (*DLEU7, MSRB3, FTO, ANAPC4, SMG6, SND1, SIM1, KCNQ5, CEP120, MACF1, SNAP91 and BCL2)* previously associated with musculoskeletal traits such as height and heel bone mineral density (Supplementary File 1 and Supplementary File 2).^24–27^ Genome-wide gene-based association analysis identified 179 genes associated with snoring beyond genome wide significance (p<2.636e-6; bonferroni corrected threshold for 18,971 tested genes) several of which were consistent with the mapped genes. After adjusting for BMI, 104 protein coding genes were identified mapping to a genome-wide significant SNP from the positional and eQTL mapping while 120 genes remained significantly associated with snoring, including both *MSRB3* and *DLEU7* (See Supplementary Figure 2 and Supplementary Files 2 and 3). eQTL data obtained from GTEx and mapped with FUMA highlighted significant SNPs that were associated with the expression of genes in several tissues including lungs, blood, oesophagus, breast mammary, tibial nerve, and several areas of the brain, such as the cerebellum and hippocampus (Supplementary Figure 3 and Supplementary File 2). In summary, many of the mapped genes for snoring have been previously associated with other traits and diseases, primarily grouped into cardiometabolic, cognitive/neurological, respiratory and psychiatric (Figure 4 and Supplementary File 1).

**Figure 4.**
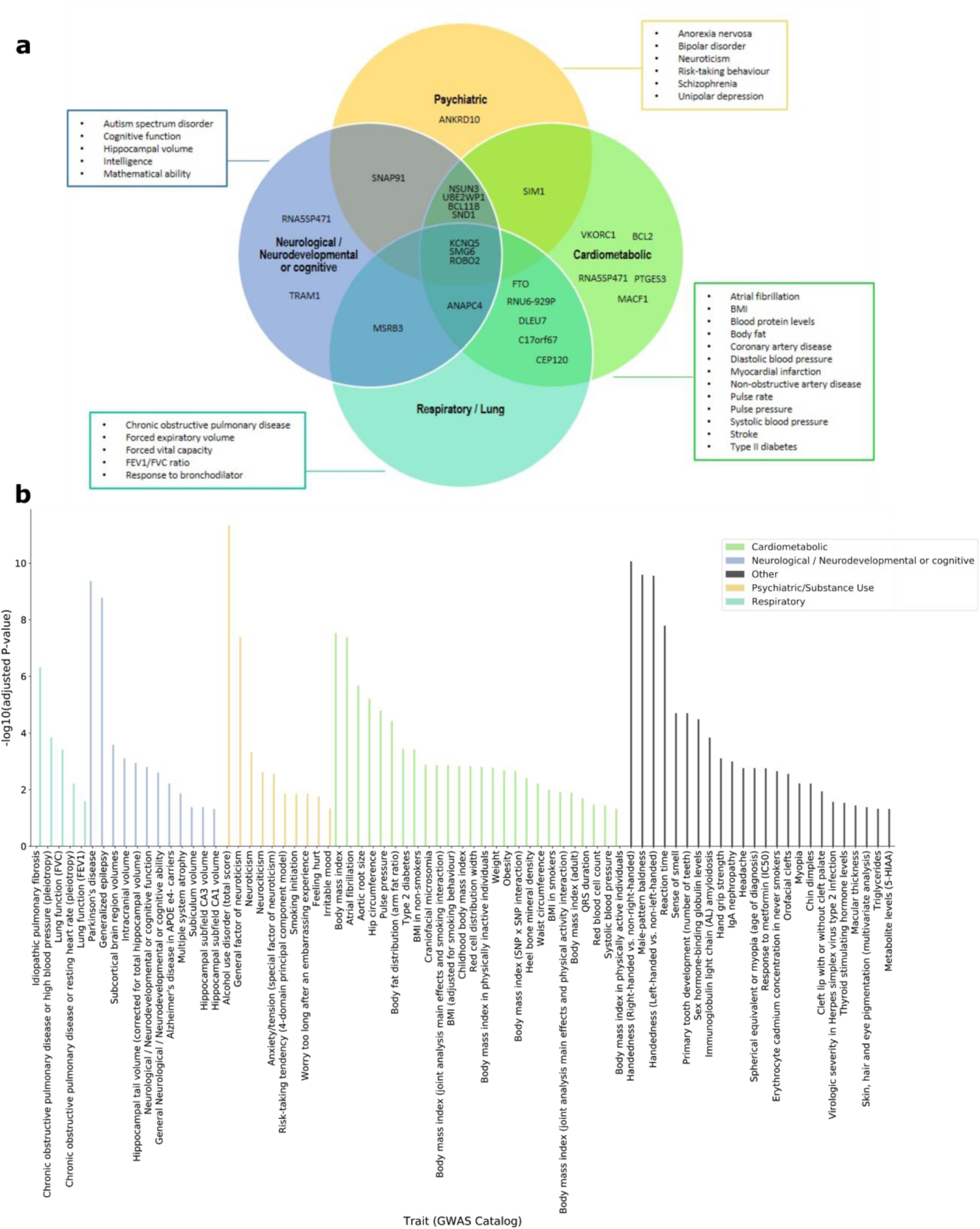
Snoring genes associated with other traits or diseases. a) Venn diagram showing nearest genes to the lead significant SNP per genomic risk loci identified for snoring, categorised according to previously reported association with other traits or diseases in the GWAS catalog. b) Significant gene set enrichment analysis (hypergeometric test) based on all prioritised genes against gene-sets defined by traits in the GWAS catalog.

To further assess whether significant genes converged in functional gene sets and pathways, we conducted gene-set enrichment analyses of tissue expression data (Supplementary Figure 3a). Genes expressed in blood vessel and tibial artery tissue were associated with snoring, even after adjusting for BMI (Supplementary Figure 3b). Given these associations, and an observed genetic correlation between snoring and pulse rate (*r*_G_= 0.106, S.E.= 0.03, p-value=0.001), we conducted two sample Generalised Summary-data-based Mendelian Randomisation (GSMR)^28^ to test for a possible causal relationship. The analysis suggested a one-way causal relationship in which snoring increased pulse rate (Supplementary Figure 4). We further explored the association between snoring and BMI or whole body fat mass. GSMR results suggested a bidirectional causal relationship, with snoring exerting a causal effect on BMI, but also BMI exerting a causal effect on snoring. However, a one-way causal relationship was seen for whole body fat mass causing snoring (Supplementary Figure 4). To control for possible confounding due to sample overlap, we repeated the Mendelian randomisation analyses using sex-stratified GWAS results (see <u>Methods</u>). This confirmed the causal associations between BMI or whole body fat mass causing snoring, while all other associations did not retain statistical significance after controlling for multiple testing (Figure 5 and Supplementary Table 2).

**Figure 5.**
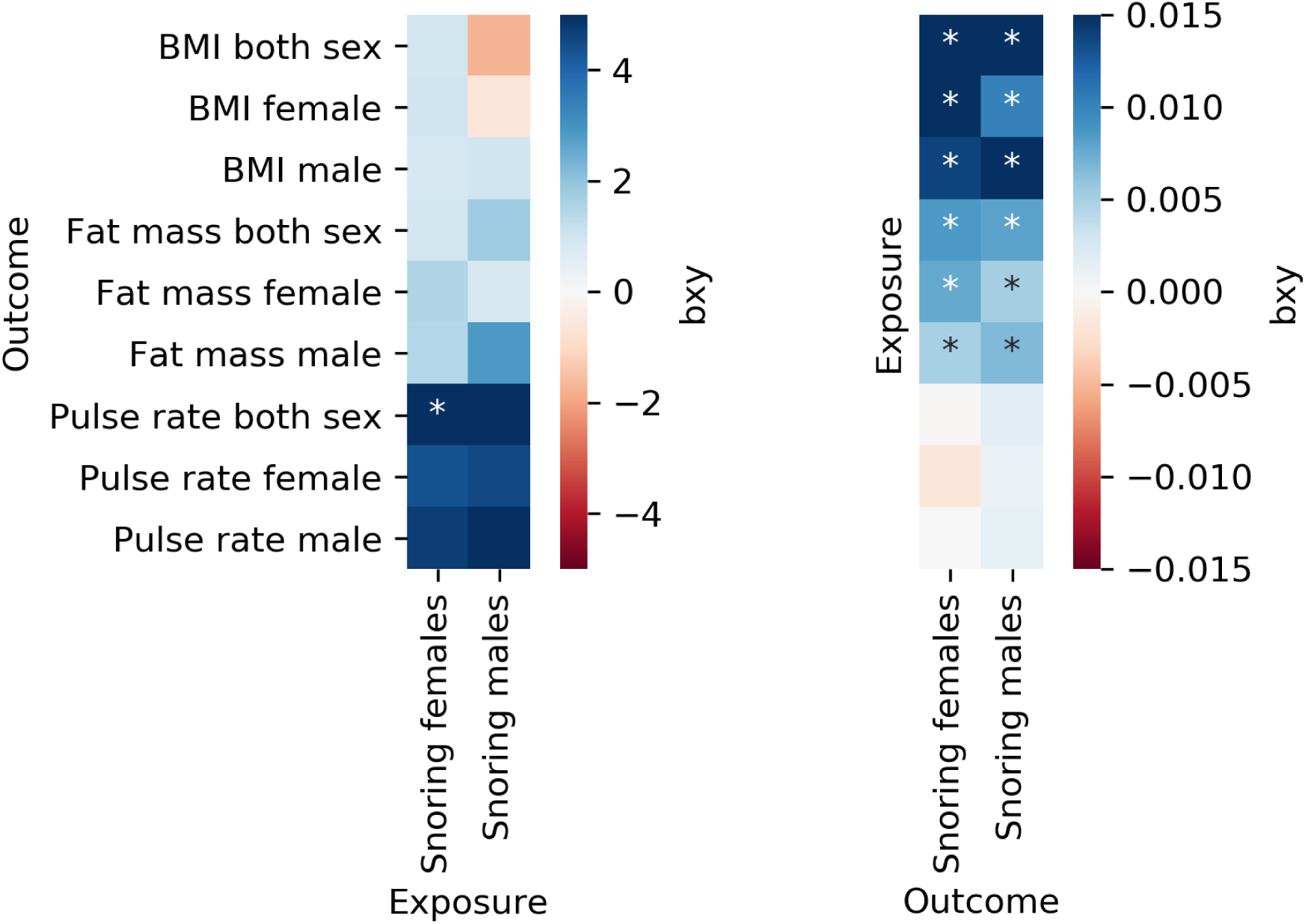
Mendelian randomisation suggests a one way causal relationship of BMI on snoring. Heatmaps depicting Mendelian randomisation results testing for causal relationships between snoring and correlated variables. Colors represent the effect size and direction as estimated using GSMR. * denotes statistical significance after Bonferroni correction for the total number of MR tests performed (p<=1.39e-03).

### Sex-stratified GWAS

Given the higher prevalence of snoring in males, we conducted GWAS analyses stratified by sex. These analyses identified 4 and 25 genome-wide significant SNPs for snoring in males and females, respectively. SNP heritability on the liability scale (*h*^*2*^_*SNP*_) was 8.77% (S.E.=0.54%) and 12.42% (S.E.=0.57%) respectively for males and females (Table 3). In the sensitivity analyses, SNP heritability (*h*^*2*^_*SNP*_) was slightly lower after adjusting for BMI in both males 7.72% (S.E.=0.56%) and females 10.85% (S.E.=0.54%) (Table 3). The cross-sex genetic correlation was high (*r*_G_=0.914, S.E.=0.033, p-value=1.91×10^−160^) and effect sizes and directions for top hits were highly consistent in both the male and female samples (Supplementary File 1 and Supplementary Figure 5).

### Polygenic scores and prediction on an independent target sample

We used the discovery GWAS summary statistics to derive PGS in an independent target sample from the Australian Genetics of Depression Study (AGDS). The prevalence of self-reported recent snoring was 32%, with a higher prevalence in males than females (43.2% and 28.1% respectively). PGS for snoring were significantly associated with recent snoring for all but one (p<=5e-8) of the p-value inclusion thresholds (Figure 6 and Supplementary Figure 6). Participants in the highest snoring PGS decile had around twice the odds of reporting recent snoring and choking or struggling for breath during sleep (i.e. probable OSA; sample prevalence=8.2%) compared to those on the lowest decile (Figure 6a). Furthermore, the PGS showed a stronger association with increasing frequency of snoring severity (Figure 6b). Finally, we showed that the snoring PGS explained a significant amount of variance in recent snoring (Figure 6c).

**Figure 6.**
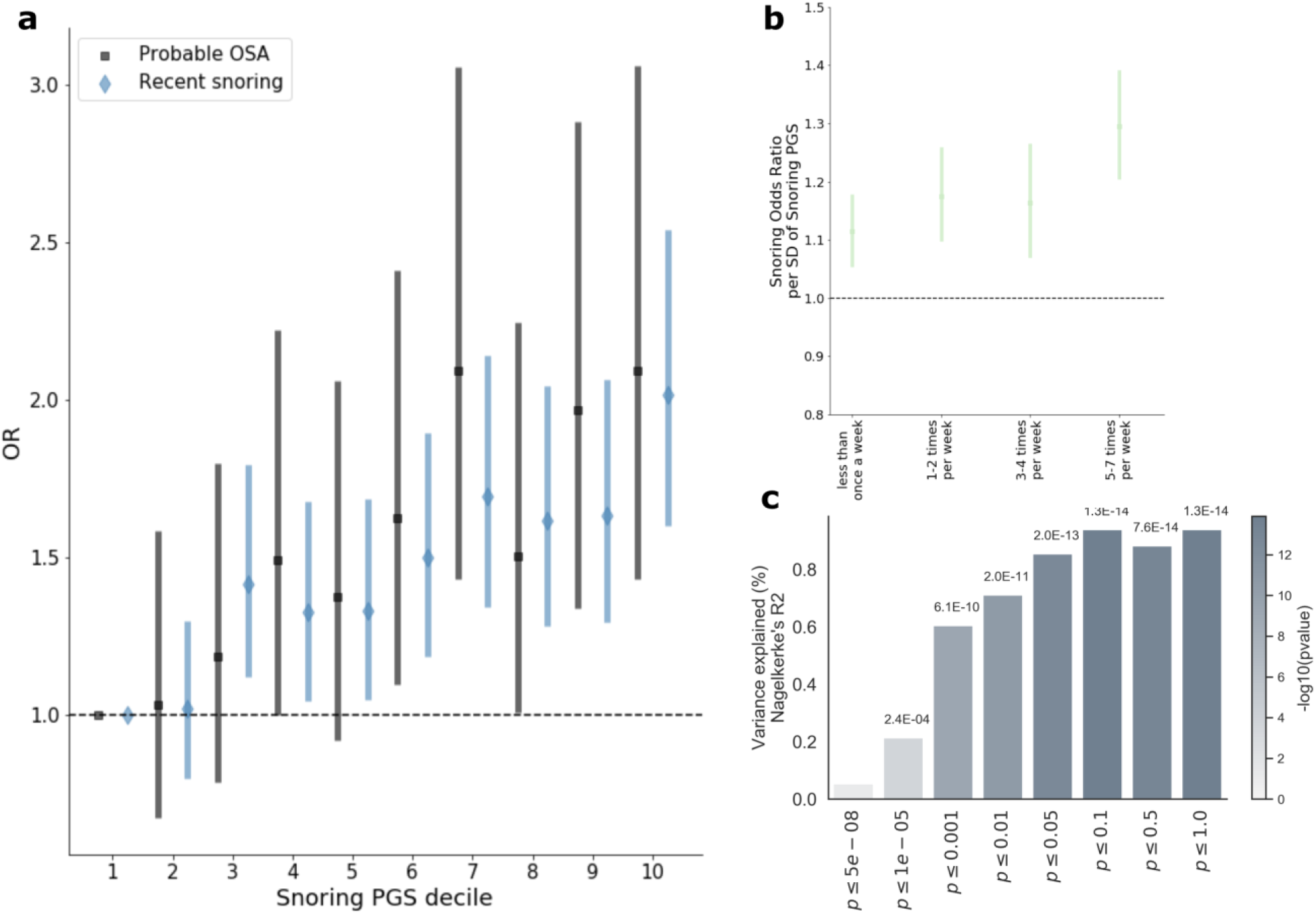
Polygenic scores predict snoring and probable apnoea in an independent sample. a) Forest plot showing the odds ratios (and 95% CI) by decile of polygenic score (PGS) for snoring from the UK Biobank discovery sample (relative to the bottom decile = 1) for recent snoring and probable sleep apnoea measured in an independent target sample of ∼8,000 unrelated Australian adults from the Australian Genetics of Depression Study (AGDS). b) Snoring PGS predicting different snoring frequencies reported in AGDS target sample. c) Variance of recent snoring in AGDS explained by PGS calculated from UK Biobank summary statistics. The x axis represents the p-value threshold used for variant inclusion during genetic scoring; the y axis represents the amount of variance explained (change in Nagelkerke R^2^). The color of each bar represents the significance of the association between the PGS and recent snoring (-log10 p-value) while the exact p-value is shown above each bar.

## DISCUSSION

This study advances our understanding of the aetiology and genetic architecture of snoring. Overall, snoring prevalence was higher in males than in females, having a strong and positive correlation with BMI, tobacco smoking and alcohol consumption in both sexes. The effects of BMI, smoking and alcohol consumption have been previously reported in other studies.^7,29,30^ In our study, tobacco smoking displayed a stronger association with snoring in females compared to males, a result consistent with an observations in an independent sample of ∼15,000 Europeans published in 2004.^30^ Previous studies provided evidence that women might have a greater susceptibility to chronic obstructive pulmonary disease after smoking^31^ and a greater bronchial hyperresponsiveness after methacholine challenge.^32^ These results suggest that smoking might be associated with snoring through an increased inflammatory response and irritation of the airways, thus having a larger effect size on females, compared to males. Conversely, our study indicates that the frequency of alcohol consumption has a stronger influence on snoring in males, compared to females. This is consistent with a previous study which failed to identify an association between alcohol consumption and snoring in females but did so in males.^33^ Our results strengthen the evidence pointing to alcohol as a risk factor for snoring and sleep apnoea, potentially through a weakening (relaxation) effect in the jaw and pharyngeal muscles.^34^ Future studies should leverage statistical genetics methods such as polygenic scoring or Mendelian randomisation to further characterise the role of smoking and alcohol-related phenotypes in snoring.

The sex differences described above motivated us to perform sex-stratified GWAS. The larger sample size of the female subgroup conferred more power to detect genetic associations in our analyses. The observed high cross-sex genetic correlation, and a high concordance in effect size and direction amongst the top hits suggests that differences in sex-stratified GWAS might be simply due to power differences between the male and female subsamples rather than the existence of sex-specific genetic effects.

Top genes identified from gene-based test analysis for snoring included *DLEU7* and *MSRB3*. Previous reports have associated *DLEU7* with heel bone mineral density,^24,35^ BMI,^36,37^ height,^38,39^ cardiovascular diseases,^40^ systolic blood pressure^40^ and pulmonary function decline (forced expiratory volume).^41^ The association between snoring genes and heel bone mineral density could be mediated by BMI due to the association between BMI and bone density documented previously.^42^ *MSRB3* plays a relevant role in protein and lipid metabolism pathways^43^ and has been associated with hippocampal volume,^44–46^ lung function,^40,47,48^ Alzheimer’s disease,^49,50^ brain injuries,^51^ novelty seeking,^52^ deafness^53^ and height.^40^ These results could be consistent with the fact that severe snoring may incur in nocturnal oxygen desaturation,^54^ diminishing neuropsychological functions and in some cases, resulting in tissue damage^55^ and contributing to impairment of memory consolidation processes.^56^ However, more research is needed in order to test this hypothesis.

Genetic correlations between snoring and a variety of traits and diseases were identified. The trait with the highest genetic correlation with snoring is sleep apnoea, which is consistent with loud snoring being a diagnostic criterion for OSA. This observation is also remarkable given the small sample size (less than 2000 cases) and therefore reduced power (no genome-wide hits) that the GWAS for self-reported sleep apnoea has in the UK Biobank. Our analyses suggest that a GWAS for snoring captures a substantial portion of the genetic contribution to sleep apnoea, highlighting the importance of studying symptoms on a subclinical threshold, an approach that has already proven useful at understanding the heterogeneity of other complex traits such as depression and neuroticism.^57,58^ Our study will enable future efforts aimed at understanding the underlying genetic architecture of OSA using multivariate statistical genetic approaches.

We also observed moderate correlations with BMI, obesity and whole body fat mass. Other relevant correlations included lung function, neurological, cardiovascular and psychiatric diseases. The high genetic correlation between snoring and snoring adj. BMI (*r*_G_=0.923, S.E.=0.003, p-value=1×10^−300^) supports the idea that the genetic architecture of snoring cannot be explained simply by BMI. This highlights the importance of studying snoring as a trait in its own, which might open new opportunities to the understanding of highly related sleeping disorders, including OSA, as well as its relationship with other traits and diseases.

Our initial Mendelian randomisation results suggested a mutual causal relationship between BMI and snoring, but only a one-way causal relationship of whole body fat mass causing snoring. We hypothesise BMI to be more heterogeneous and potentially more pleiotropic than whole body fat mass. In fact Mendelian randomisation is known to be confounded by pleiotropy.^59^ Interestingly, a one-way causal relationship between snoring and pulse rate, which survived adjustment for BMI, was identified. Nonetheless, this association did not reach statistical significance when accounting for sample overlap (i.e. sex-stratified GSMR). The only causal associations that survived this were BMI or whole body fat mass causing snoring. The significantly lower number of instruments available for snoring as an exposure (Supplementary Table 2) makes it hard to assess whether the results were due to a lack of power or a lack of true causal effect. Future efforts could leverage novel statistical genetics methods that use all the GWAS results to test whether the associations observed could be explained by a causality rather than pleiotropy.^60^

Finally, we assessed the validity of our GWAS results by using genetic scoring on an independent sample of Australian adults with data on *recent snoring*. Our successful prediction of snoring using PGS supports the external validity of our genetic association results. Remarkably, we predicted *probable OSA* using a snoring derived PGS. Thus, investigating the aetiology of snoring could also help uncover the aetiology and genetic architecture of OSA, a task that has proved difficult and challenging^61^. Future efforts could assess whether the utility of snoring derived PGS as an addition to the current battery of tests used to more accurately diagnose OSA,^62^ particularly given the issue of potential OSA underdiagnosis.^63,64^

Our results highlight the relevance of studying snoring as an independent trait, and provide important insights into its aetiology and genetic architecture. This study represents the largest genetic investigation of snoring to date. However, some limitations must be acknowledged. Analyses used self-reported snoring with the item relying on a partner or close friend complaining about the participant’s snoring. Thus, the case definition might be subject to participant specific recall and subjective biases. Nonetheless, we hypothesize that this limitation might result in the inclusion of some cases as controls (i.e. snoring participants living alone) and therefore bias our results toward the null rather than creating false positives. To avoid confounding due to population stratification, we only included samples of European ancestry in our analyses. This is particularly important given reports of ethnic differences associated with snoring prevalences.^65^ Nonetheless, excluding other populations can limit the generalisability of these results outside the populations studied. As previously discussed, we cannot identify which, if any, sex-specific genetic observations (e.g. differences in SNP heritability) are due to true genetic effects rather than power differences between the samples. Finally, the fact that PGS for snoring predicted less than one percent of the variance on recent snoring suggest that the GWAS is still underpowered^66^. The heritability for snoring in twin studies is estimated in the range of 18 to 28%^6^, although some of the *missing heritability* for snoring may come from dominant genetic effects, it is likely that an increased power and studying rare variants^67^ yields more powered genetic predictors.

## CONCLUSIONS

In summary, we provide insights into the aetiology of snoring, its risk factors and genetic underpinnings. Our observational analyses showed a higher prevalence of snoring in males compared to females, and effects of age, BMI, smoking and alcohol consumption. In addition, tobacco smoking showed a higher effect on snoring prevalence in females compared to males while alcohol consumption displayed a higher effect on snoring prevalence in males compared to females. GWAS identified 127 genome-wide significant associations across 41 genomic risk loci with *h*^*2*^_*SNP*_ = 9.9%. We found no evidence for sex-specific genetic factors, and showed that most of the SNP heritability identified is not simply due to BMI. We also found evidence of a causal relationship from BMI or whole body fat mass to snoring. We further found evidence of genetic overlap between snoring and other cardiometabolic, respiratory, neurological and psychiatric traits. Finally, we used the GWAS summary statistics to derive individual PGS and predict both recent snoring and probable OSA in an independent sample of Australian adults, thus confirming the relevance of snoring as a sleep-related complex trait. Future studies should aim at leveraging powered GWAS on alcohol and tobacco behaviours to assess whether they are truly causal of snoring, and to assess the amount of shared genetic overlap between OSA and habitual snoring, as the latter may serve to boost the power of obstructive sleep apnoea genetic studies.

## METHODS

### Discovery sample and phenotypic information

Participants included in the present study were of European ancestry from the UK Biobank. Briefly, this resource recruited participants between 2006 and 2010 to assess lifestyle, anthropometric and health related variables. Participants self-reported on sleep-related traits. Snoring was assessed as a single item (Field-ID: 1210): “Does your partner or a close relative or friend complain about your snoring?” This question could be answered with “Yes”, “No”, “Don’t know”, or “Prefer not to answer”. We excluded participants whose answers were “Don’t know” (*n* = 29,309) or “Prefer not to answer” (*n* = 6,854) from our analyses. (Supplementary Table 3 shows the total sample size for each GWAS, including sensitivity and sex-stratified analyses). OSA cases were determined on the basis of either ICD-10 diagnosis code or self-report of *sleep apnoea* diagnosis in the UK Biobank.

### Data extraction and statistical analyses

Raw data was extracted from the UK Biobank under Application Number 25331. For a description of the field codes and instances used refer to Supplementary Table 4. Data was re-coded to remove missing data and uninformative responses (e.g. “I don’t know” or “I would rather not answer”). Phenotype derived estimates such as prevalence and associations between variables were calculated using *python*. Libraries such as *numpy* (https://docs.scipy.org/doc/numpy/user/) and *scipy* (https://docs.scipy.org/doc/) were used for descriptive statistics and *statsmodels* (http://conference.scipy.org/proceedings/scipy2010/pdfs/seabold.pdf) was used to build logistic regression models to assess correlates of snoring and calculate odds ratios. Snoring prevalence stratified plots (e.g. Figure 1) were performed using *seaborn* v0.9.0. Confidence intervals were calculated using a bootstrapping procedure generating 1000 pseudo-replicates of the data.

### Genetic association analyses and quality control

#### Discovery GWAS

Analyses were performed for both sex stratified and full sample using the BOLT-LMM software tool. We used a strict quality control (QC) procedure corresponding to minor allele frequency (MAF>=0.005) and imputation quality (>=0.60).

#### Sensitivity analyses

Given the strong correlation between snoring and BMI, we carried out GWAS for snoring using BMI as a covariate (snoring adj. BMI) with BOLT-LMM software with a quality control corresponding to minor allele frequency (MAF>=0.005) and imputation quality (>=0.60).

### Post-GWAS Annotation and Functional Mapping

SNP annotation was conducted using the FUMA platform. Risk loci are defined as up to 250 kb based on the most right and left SNPs from each LD block. Gene based tests were performed using Multi-marker Analysis of GenoMic Annotation (MAGMA) as implemented on the FUMA platform, which provides aggregate association p-values based on all variants located within a gene and its regulatory region. We used the GWAS summary statistics to conduct a MAGMA^68^ analysis in the FUMA^13^ platform (https://fuma.ctglab.nl/). This analysis includes a gene-based test to detect significant SNPs associated with snoring. The prioritised genes based on positional and eQTL mapping were further used to perform gene-set enrichment analysis against the traits available in the GWAS catalog. Furthermore, we used FUMA to perform tissue enrichment analysis, based on data from the Genotype-Tissue Expression (GTEx) project (https://gtexportal.org/home/documentationPage).

### Genetic correlation analyses

We performed genetic correlation analyses to estimate genetic correlations between the discovery, sensitivity and sex-stratified snoring GWAS summary statistics using LD score regression as implemented in the Complex Trait Genomics Virtual Lab (CTG-VL, http://genoma.io). Further, in order to uncover genetically correlated traits with snoring, genetic correlation analyses using LD Score Regression (LDSC) were performed on the platforms CTG-VL and LDHub (http://ldsc.broadinstitute.org/ldhub/), which aggregate summary statistics for GWAS on hundreds of traits.

### Mendelian randomisation

Mendelian randomisation (MR) is a method in which genetic variants (e.g. single nucleotide polymorphisms, SNPs) are used as instrumental variables to determine causal relationships between traits, environmental exposures, biomarkers, or disease outcomes^69^; to satisfy the conditions for MR, it is not required to identify the actual causal variant because a marker in Linkage Disequilibrium (LD) with the causal variant can serve as a proxy instrument^70^. Moreover, in order to draw conclusions in regard with casual effects, three relevant assumptions must be taken into consideration: (I) genetic variants must be associated with the exposure of interest; (II) genetic variants must not be confounded; (III) genetic variants must be independent of the outcome through other mechanisms.^71^ We used Generalised Summary-data-based Mendelian Randomisation (GSMR)^70^, an approach that leverages the usage of multiple independent variables (SNPs) strongly associated with the outcome to overcome these assumptions, as implemented in the CTG -VL (http://genoma.io). We used GSMR to assess causal relationships between snoring and: BMI, whole body fat mass and pulse rate using our results and existing summary statistics for these traits. To avoid possible confounding from sample overlap, we performed GSMR using the summary statistics derived from sex-stratified GWAS. For example, the female snoring GWAS results were used as exposure while the male pulse rate GWAS results were used for the outcome.

### Target sample and polygenic scoring

In order to quantify for the cumulative genetic associations for snoring, we calculated PGS using a *clumping* + *thresholding* approach. Study description and sample characteristics of the Australian Genetics of Depression study is available elsewhere.^72^ Genotyping was conducted using the Illumina Infinium Global Screening Array platform and genotype imputation using the Haplotype Reference Consortium’s reference panel in the Michigan Imputation Server^73^ was carried out after performing standard quality control procedures. Briefly, for PGS estimation, we excluded indel, strand ambiguous- and low (R2<0.6) imputation quality-variants. The most significant SNPs were selected using a conservative clumping procedure (PLINK1.9^74^; p1=1, p2=1, r2=0.1, kb=10000) to correct for inflation arising from linkage disequilibrium (LD). Eight PGS were calculated using different p-value thresholds (p<5×10-8, p<1×10-5, p<0.001, p<0.01, p<0.05, p<0.1, p<0.5, p<1) as criteria for SNP inclusion on the PGS calculation. PGS were calculated by multiplying the effect size of a given SNP by the imputed number of copies (using dosage probabilities) of the effect allele present in an individual. Finally, the SNP dosage effects were summed across all loci per individual. To assess the association between the PGS and snoring and probable OSA, we employed a logistic regression (python *statsmodels*). The target sample for snoring was a subset (n=9,026) of the Australian Genetics of Depression Study (AGDS) with data on recent snoring collected through the self-reported item: “*During the last month, on how many nights or days per week have you had or been told you had loud snoring*”. The item for probable OSA was: “*During the last month, on how many nights or days per week have you had or been told your breathing stops or you choke or struggle for breath*”. For both items a positive response was considered from *1-2 times per week* up to *5-7 times per week* and the answer *Rarely, less than once a week* was excluded. Only a subset (n∼8,000) of highly unrelated individuals (genetic relatedness <0.05) were included in the analyses.

## ACKNOWLEDGEMENTS

This research was conducted using data from the UK Biobank resource (application number 25331). Data collection for the Australian Genetics of Depression Study was possible thanks to funding from the Australian National Health & Medical Research Council (NHMRC) to NGM (GNT1086683). AC-G is supported by a UQ Research Training Scholarship from The University of Queensland (UQ). MER thanks the support of the NHMRC and Australian Research Council (ARC), through a NHMRC-ARC Dementia Research Development Fellowship (GNT1102821). GC-P is funded by an ARC Discovery Early Career Researcher Award (DE180100976).

## AUTHOR CONTRIBUTIONS

MER and GC-P conceived and directed the study. AIC and LMG-M performed most of the statistical and bioinformatics analyses, with support from GC-P and MER. EMB and NGM collected and contributed data from the Australian sample. AIC, LMG-M, MER and GC-P wrote the paper with feedback from all co-authors.

## Supplementary file description

**Supplementary Materials**. Supplementary Figures 1-6 and Supplementary Tables 1 - 4

**Supplementary File 1**. Genomic risk loci for discovery, sensitivity and sex-stratified analyses.

**Supplementary File 2**. FUMA output files for snoring and genetic correlations.

**Supplementary File 3**. FUMA output files for snoring adjusted for BMI and genetic correlations.

