## Supplementary Figure for "Insights into the aetiology of snoring from observational and genetic investigations in the UK Biobank (n=408,317)"

**SUPPLEMENTARY FIGURES AND TABLES**

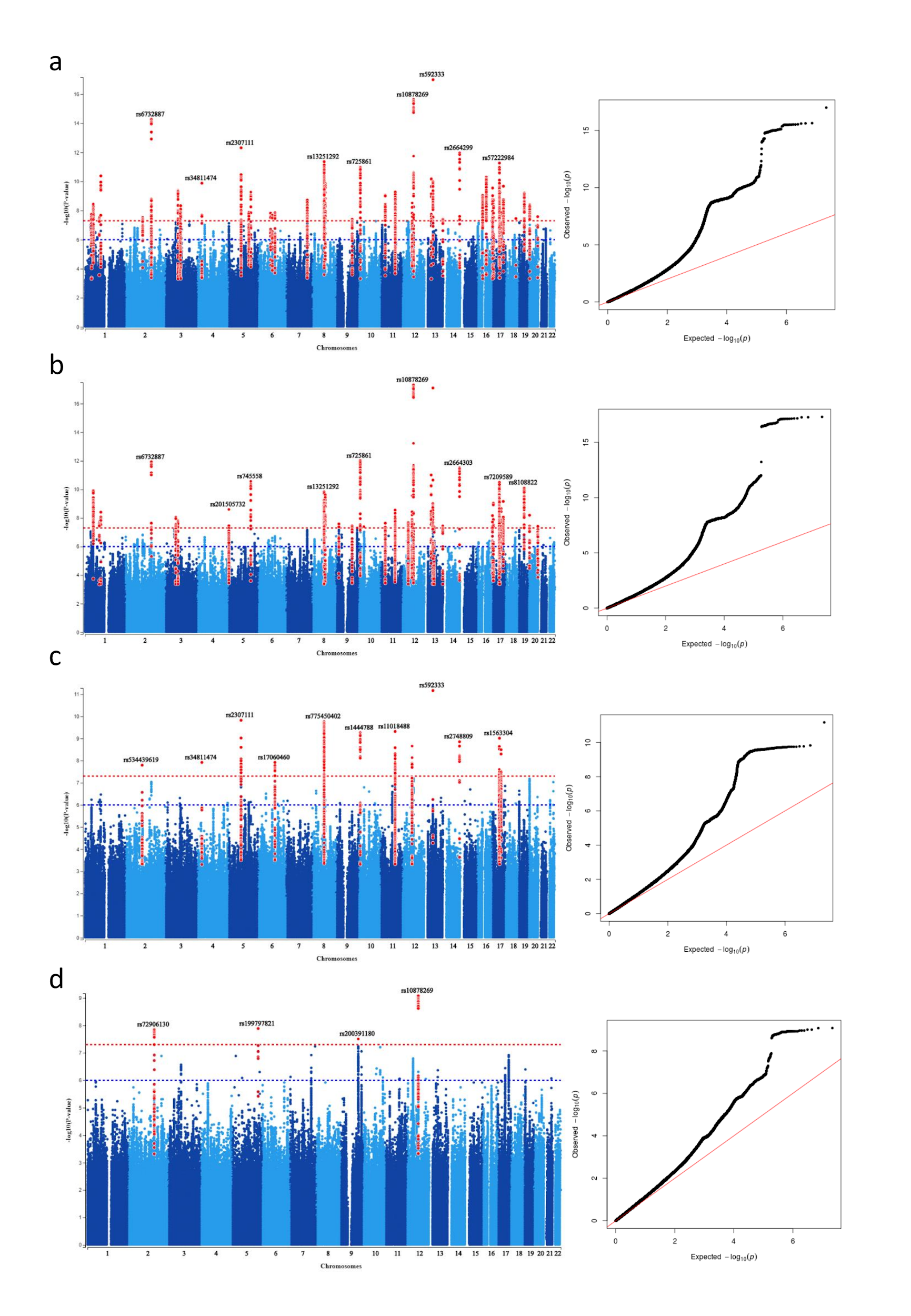

**Supplementary Figure 1. GWAS and sensitivity GWAS results.**

Supplementary annotated Manhattan plots with LD highlighted and top hits labelled (left panels). quantile-quantile plots (QQplots) are depicted on the right panels. a) Results for snoring GWAS. b) Results for snoring GWAS adjusting for BMI as a covariate. c) Results for snoring GWAS performed only on the female subset of the sample. d) Results for snoring GWAS performed only on the male subset of the sample.

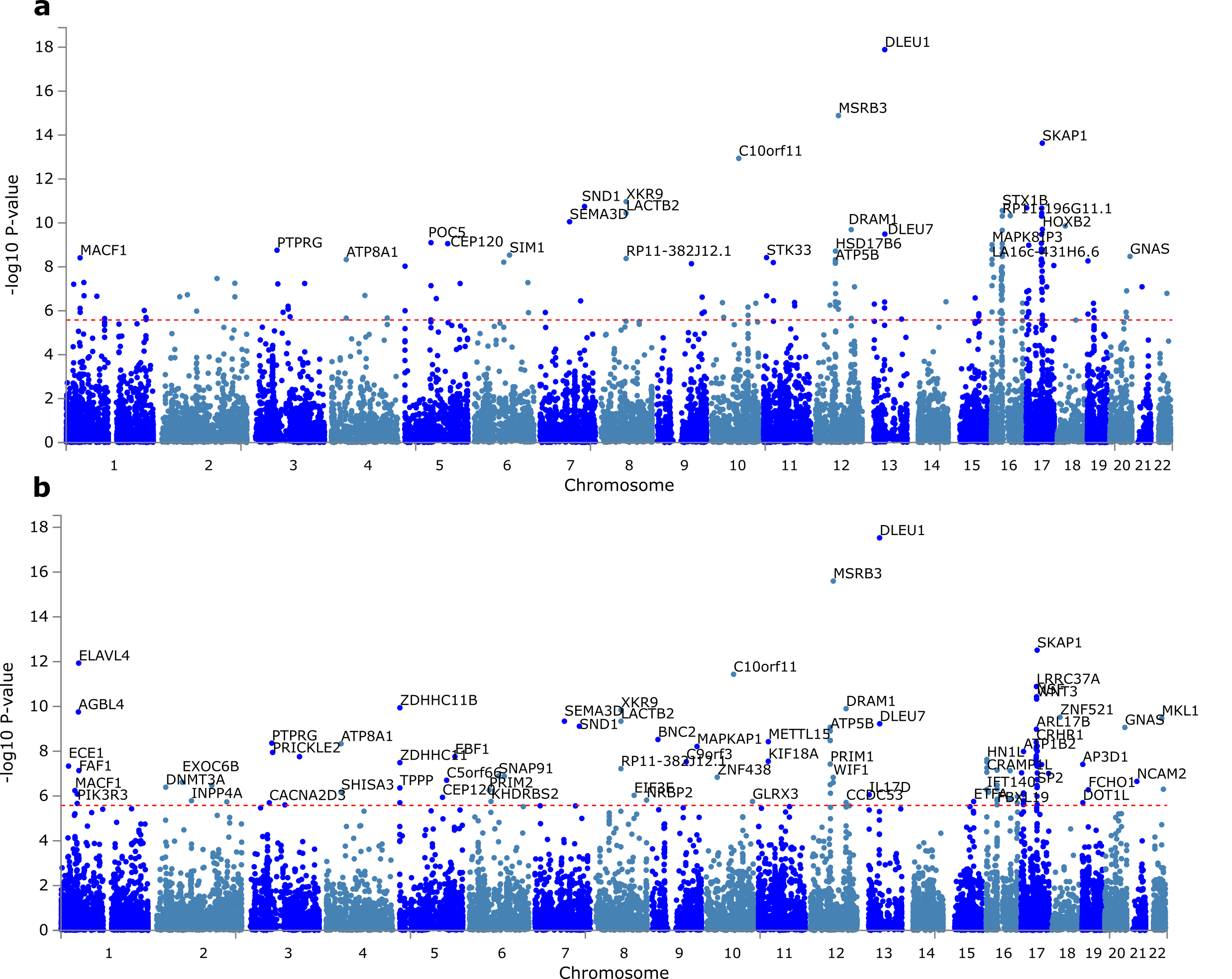

**Supplementary Figure 2. MAGMA Gene-based test association analyses**

Manhattan plots depicting the gene-based test association analyses performed with MAGMA for snoring (a) and snoring adjusted for BMI (b). Some gene labels were removed or rearranged for better clarity. The red line indicated the bonferroni corrected genome-wide significance threshold (p<2.636e-6; 18,971 tested genes).

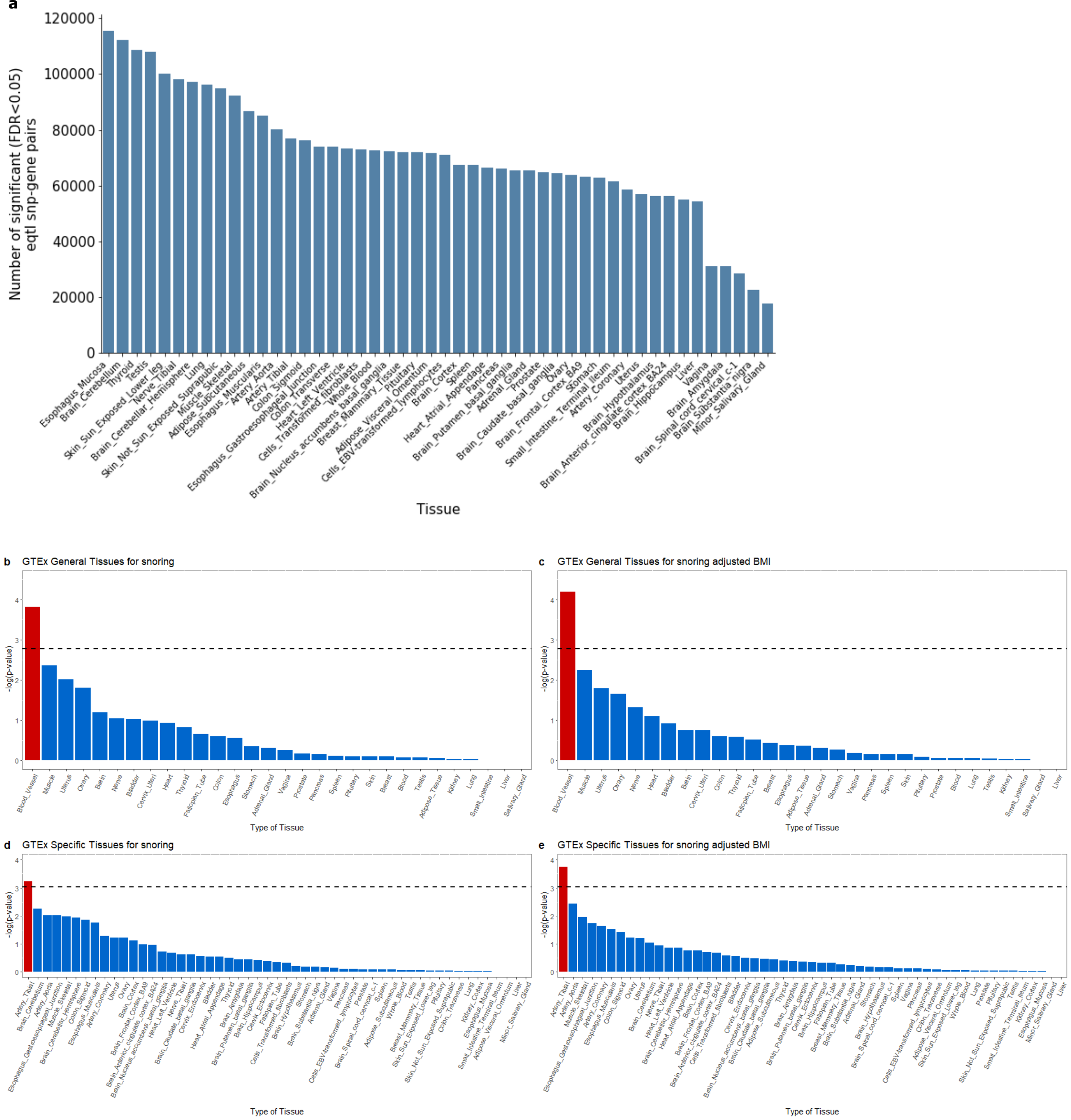

**Supplementary Figure 3. MAGMA Tissue Enrichment Analysis.**

Bar plots showing gene set enrichment analyses for eQTL mapping (a) and tissue-specific gene expression based on the GWAS for snoring (a,b,d) and snoring adjusted for BMI (c,e). Analyses performed with MAGMA. Red bars indicate significantly enriched tissues (i.e. crossing the bonferroni corrected significance threshold shown with a dashed line).

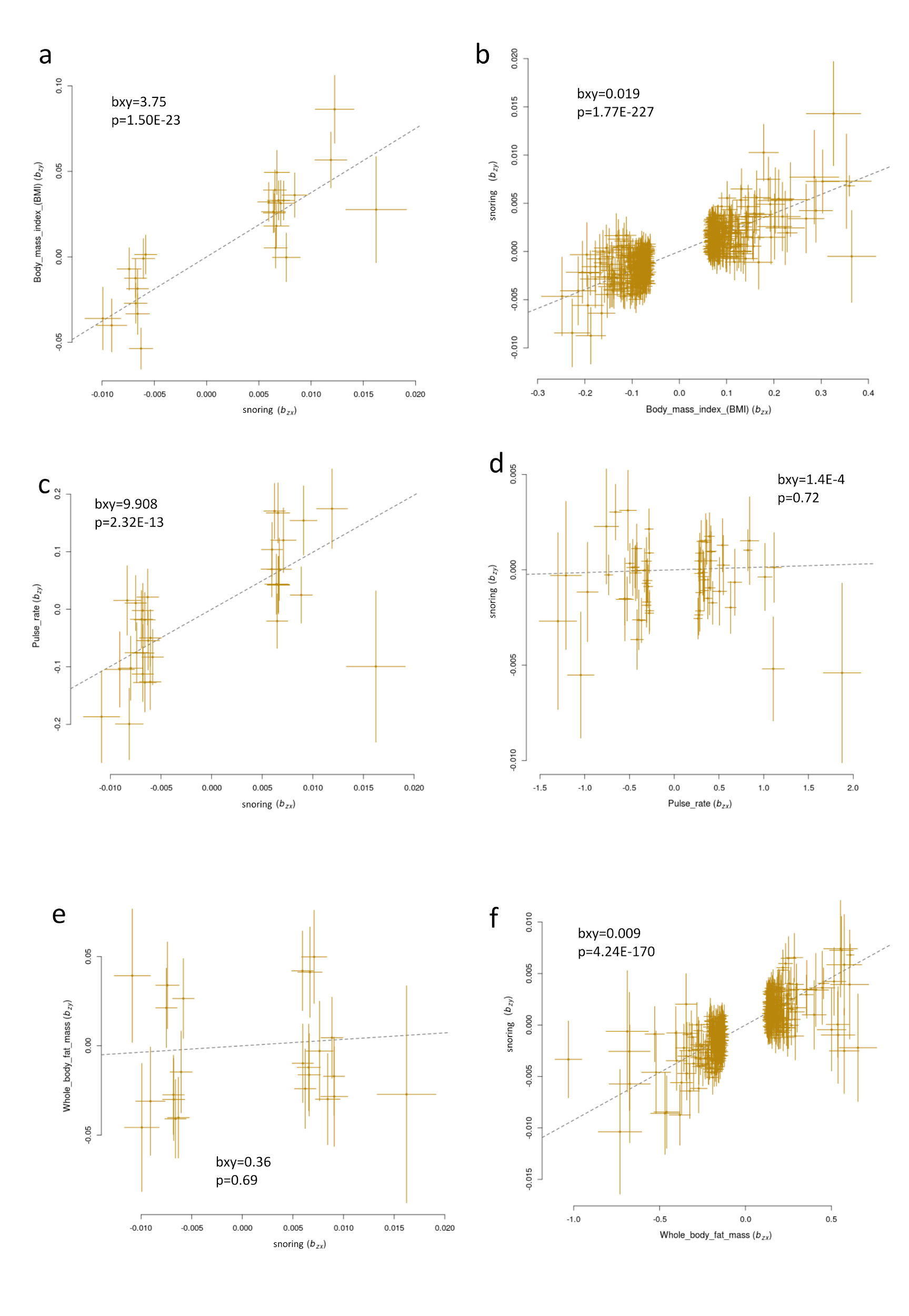

**Supplementary Figure 4. Mendelian randomisation effect size plots**

Scatter plots depicting the relationship between the effect sizes of SNP instruments for exposures (x-axis) and outcomes (y-axis). If the exposure and the outcome have a causal relationship, then the effects of SNPs (instruments) on the exposure should have proportional effects on trait two. Inset of each plot depicts the estimated causal effect (bxy) and its associated p-value. a,c,e) Show the results of snoring as an exposure while b,d and f show the effects of BMI, Pulse rate or Whole body fat mass as exposures.

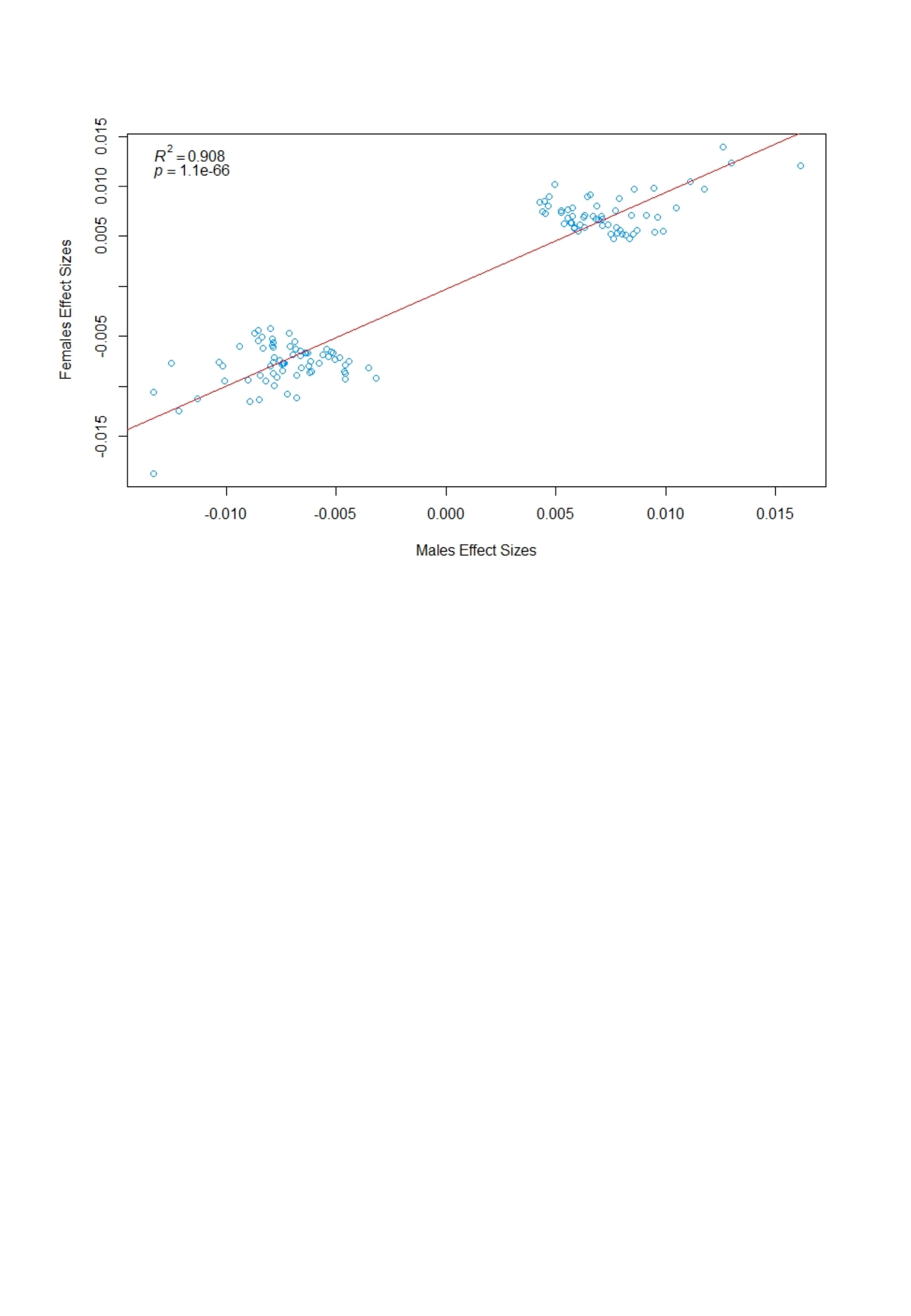

**Supplementary Figure 5. Cross-sex effects size of snoring associated SNPs.**

Scatter plot showing the effect sizes of the main GWAS independent SNPs across males (x-axis) and females (y-axis). The line represents a linear regression showing a high concordance between sex association results.

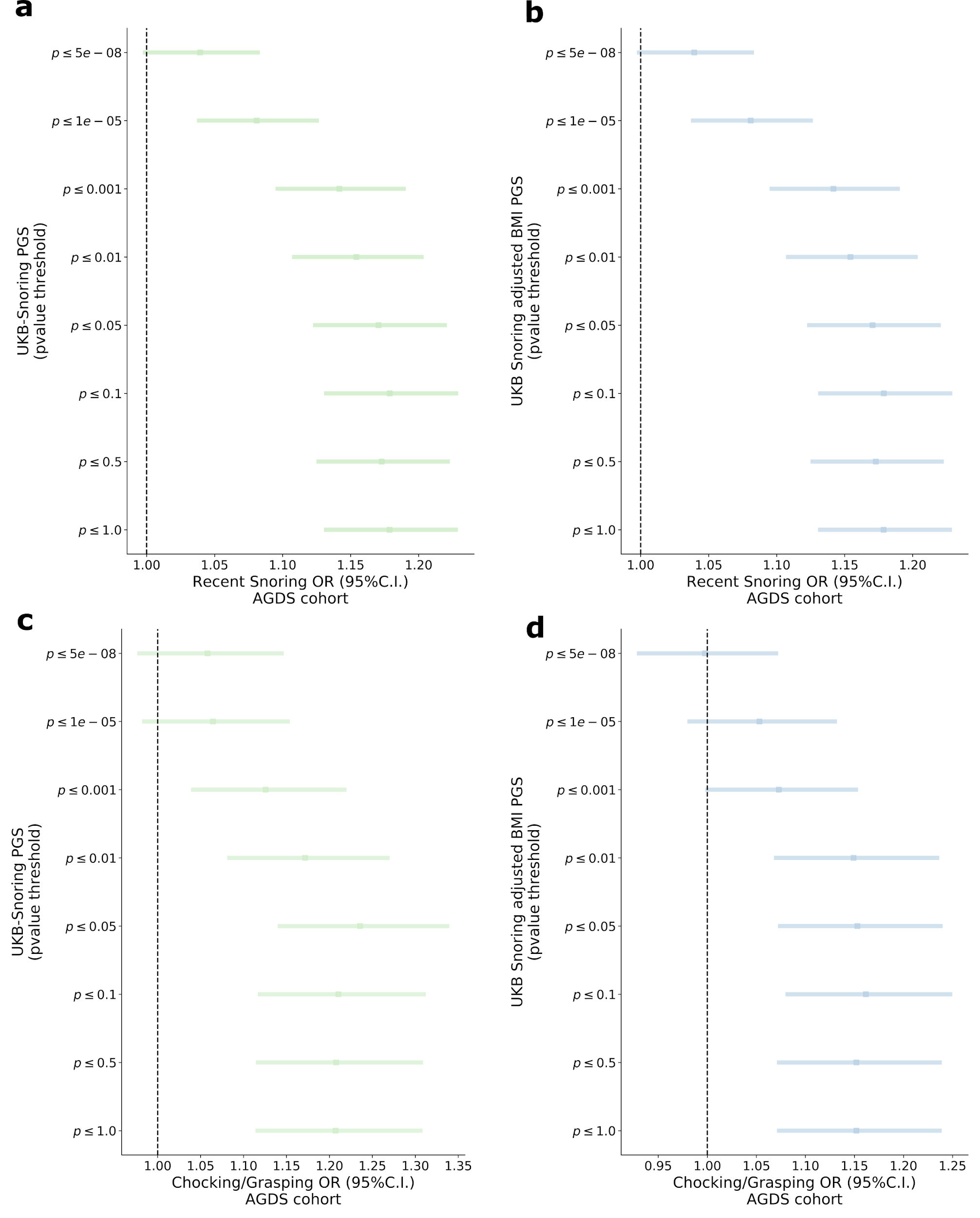

**Supplementary Figure 6. Genetic scoring results**

Results of association between snoring (a,b) or snoring adjusted for BMI (c,d) and polygenic scores (PGS) for snoring constructed with an increasingly stringent p-value threshold for variant inclusion (y axis). The x-axis denotes the odds ratio (OR) between the PGS and the phenotype. Bars denote the 95% confidence interval.

| **Supplementary Table 1.** Phenotypic correlations of snoring and associated factors | | | |
| --- | --- | --- | --- |
| **Predictor** | **beta (logOR)** | **S.E.** | **P-value** |
| Intercept | -4.936 | 0.09 | 0.0E+00 |
| Current tobacco [Occasionally] | 0.1539 | 0.04 | 6.1E-05 |
| Current tobacco [Most Days] | 0.3697 | 0.03 | 3.3E-41 |
| Alcohol frequency [rarely] | 0.1031 | 0.03 | 1.5E-04 |
| Alcohol frequency [occasionally] | 0.1467 | 0.03 | 9.5E-09 |
| Alcohol frequency [frequently] | 0.2209 | 0.02 | 3.9E-20 |
| Alcohol frequency [ very frequently] | 0.319 | 0.02 | 2.3E-40 |
| Pulse rate | 0.003 | 0 | 1.0E-07 |
| Sleep_duration | 0.0547 | 0.01 | 2.0E-21 |
| Whole body fat mass | 0.0051 | 0 | 4.7E-03 |
| Sex | 0.846 | 0.02 | 0.0E+00 |
| BMI | 0.0891 | 0 | 4.0E-136 |
| Age | 0.0103 | 0 | 1.9E-39 |
| Apnoea | 1.1206 | 0.06 | 4.5E-80 |

| **Supplementary Table2** Mendelian randomisation results | | | | | |
| --- | --- | --- | --- | --- | --- |
| Exposure | Outcome | bxy | se | p | nsnp |
| Snoring males | Fat mass both sex | 1.78 | 1.33 | 0.1826 | 3 |
| Snoring males | Fat mass female | 0.85 | 1.97 | 0.6663 | 3 |
| Snoring males | Fat mass male | 2.87 | 1.76 | 0.1028 | 3 |
| Snoring males | BMI both sex | -1.73 | 0.85 | 0.0417 | 2 |
| Snoring males | BMI female | -0.65 | 1.02 | 0.5211 | 3 |
| Snoring males | BMI male | 0.99 | 0.90 | 0.2699 | 3 |
| Snoring males | Pulse rate both sex | 5.58 | 2.90 | 0.0541 | 3 |
| Snoring males | Pulse rate female | 4.51 | 3.80 | 0.2343 | 3 |
| Snoring males | Pulse rate male | 6.78 | 4.50 | 0.1319 | 3 |
| Snoring females | Fat mass both sex | 0.95 | 0.80 | 0.2347 | 11 |
| Snoring females | Fat mass female | 1.51 | 1.19 | 0.2034 | 11 |
| Snoring females | Fat mass male | 1.46 | 1.01 | 0.1496 | 12 |
| Snoring females | BMI both sex | 0.94 | 0.41 | 0.0223 | 11 |
| Snoring females | BMI female | 1.00 | 0.61 | 0.1015 | 11 |
| Snoring females | BMI male | 0.88 | 0.54 | 0.1042 | 11 |
| Snoring females | Pulse rate both sex | 5.55 | 1.72 | 0.0013 | 12 |
| Snoring females | Pulse rate female | 4.37 | 2.21 | 0.0484 | 12 |
| Snoring females | Pulse rate male | 4.76 | 2.44 | 0.0513 | 13 |
| Fat mass both sex | Snoring males | 8.00E-03 | 4.54E-04 | 2.20E-69 | 600 |
| Fat mass female | Snoring males | 5.06E-03 | 5.40E-04 | 6.7E-21 | 207 |
| Fat mass male | Snoring males | 6.64E-03 | 7.06E-04 | 5.2E-21 | 146 |
| BMI both sex | Snoring males | 1.88E-02 | 8.72E-04 | 8.6E-103 | 634 |
| BMI female | Snoring males | 1.01E-02 | 1.03E-03 | 1.2E-22 | 211 |
| BMI male | Snoring males | 1.51E-02 | 1.26E-03 | 5.3E-33 | 169 |
| Pulse rate both sex | Snoring males | 1.57E-03 | 5.67E-04 | 5.6E-03 | 71 |
| Pulse rate female | Snoring males | 1.04E-03 | 6.33E-04 | 1.0E-01 | 38 |
| Pulse rate male | Snoring males | 1.28E-03 | 7.07E-04 | 7.0E-02 | 23 |
| Fat mass both sex | Snoring females | 8.57E-03 | 3.90E-04 | 5.3E-107 | 586 |
| Fat mass female | Snoring females | 7.71E-03 | 4.80E-04 | 4.5E-58 | 195 |
| Fat mass male | Snoring females | 5.04E-03 | 5.98E-04 | 3.5E-17 | 144 |
| BMI both sex | Snoring females | 1.72E-02 | 7.27E-04 | 2.9E-124 | 633 |
| BMI female | Snoring females | 1.49E-02 | 9.19E-04 | 1.8E-59 | 198 |
| BMI male | Snoring females | 1.39E-02 | 1.10E-03 | 1.1E-36 | 159 |
| Pulse rate both sex | Snoring females | -1.92E-04 | 4.91E-04 | 7.0E-01 | 70 |
| Pulse rate female | Snoring females | -1.79E-03 | 5.65E-04 | 1.6E-03 | 35 |
| Pulse rate male | Snoring females | -4.54E-05 | 6.06E-04 | 9.4E-01 | 23 |
| Full results for the mendelian randomisation shown on Figure 5. nsnp is the number of SNPs used as instrument variables. | | | | | |

| **Supplementary Table 3.** Sample sizes for discovery, sex-stratified and sensitivity GWAS analyses. | | |
| --- | --- | --- |
| **Model** | **Sample Size*** | **Snoring prevalence** |
| Snoring | 408,317 | 37.3% |
| Snoring Males | 189,971 | 47.8% |
| Snoring Females | 218,346 | 28.3% |
| BMI adjusted | 407,066 | 37.3% |
| BMI adjusted Males | 189,333 | 47.8% |
| BMI adjusted Females | 217,733 | 28.3% |
| *Sample size for the GWAS using only individuals with non-missing phenotypic and genetic data and passing quality control ancestry filters (see methods) | | |

**Supplementary Table 4.** Field codes and instances used from UK Biobank.

| **Description** | **Field Code** | **Instance** |
| --- | --- | --- |
| Age | 21022 | Single instance |
| BMI | 21001 | 0 |
| Current tobacco smoking | 1239 | 0 |
| Frequency of drinking alcohol | 20414 | Single instance |
| How are people in household related to participant | 6141 | 0 |
| Sex | 31 | Single instance |
| Snoring | 1210 | 0 |
